# LIS1, a glyco-humanized swine polyclonal anti-lymphocyte globulin, as a novel induction treatment in solid organ transplantation

**DOI:** 10.1101/2022.09.08.507114

**Authors:** Juliette Rousse, Pierre-Joseph Royer, Gwenaëlle Evanno, Elsa Lheriteau, Carine Ciron, Apolline Salama, Françoise Shneiker, Roberto Duchi, Andrea Perota, Cesare Galli, Emmanuele Cozzi, Gilles Blancho, Odile Duvaux, Sophie Brouard, Jean-Paul Soulillou, Jean-Marie Bach, Bernard Vanhove

## Abstract

Anti-thymocyte/lymphocyte globulins (ATGs/ALGs) are immunosuppressive drugs used in induction therapies to prevent acute rejection in solid organ transplantation. Because of animal origin, ATGs/ALGs contain highly immunogenic carbohydrate xenoantigens eliciting antibodies that are associated with subclinical inflammatory events possibly impacting long-term graft survival. Their strong and long-lasting lymphodepleting activity also increases the risk for infections. To circumvent these drawbacks, LIS1 has been engineered as a glyco-humanized polyclonal antibody obtained by immunizing genetically modified pigs knocked out for major xeno-antigens αGal and Neu5Gc. It is Fc-silenced in humans and differs from other ATGs/ALGs by its mechanism of action excluding antibody-dependent cell-mediated cytotoxicity and being restricted to complement and phagocyte-mediated cytotoxicity, apoptosis and antigen masking, resulting in profound inhibition of T-cell alloreactivity in mixed-leucocyte-reactions. Preclinical evaluation in macaques showed that LIS1 impacted CD4^+^, CD8^+^ effector T cells but not T-reg, B cells or myeloid cells. Compared with rabbit ATG, LIS1 induced transient depletion of target T cells in the peripheral blood but was equivalent in preventing allograft rejection in a skin allograft model. The novel therapeutic modality of LIS1 might present advantages in induction treatment after organ transplantation by shortening the T-cell depletion period while maintaining adequate immunosuppression and reducing immunogenicity.

## 1. INTRODUCTION

“Induction therapies” in solid organ transplantation refer to early lymphodepletion or blockade to avoid acute graft rejection ^1,2^. They comprise CD25 antagonists^3^ and anti-lymphocyte/thymocyte globulins (ALGs/ATGs) ^4,5^. ALGs/ATGs are polyclonal antibodies of animal origin directed against T- and B-lymphocytes^6^, obtained from rabbits or horses immunized with primary human thymocytes or human T-cell lines ^7,8^. Their main mechanism of action is apoptosis^9,10^, complement-dependent cytotoxicity (CDC), antibody-dependent cell cytotoxicity (ADCC), phagocytosis and other nondestructive mechanisms^6^. ALGs/ATGs have been used for 50 years^4,6^ and have proven efficacy in preventing acute graft rejection. However, their strong immunogenicity might be associated with side effects ranging from mild fever or skin rashes to more serious serum sickness disease (SSD) ^11,12^ or anaphylactic shock ^13–15^.

SSD is a hypersensitivity reaction toward glycoproteins from animal sources ^16^ caused by antibodies directed against glycans bearing the common mammalian sialic acid N-glycolylneuraminic acid (Neu5Gc)^17–21^. Humans present a biased sialylation of glycoproteins and glycolipids. The lack of Neu5Gc^22^ on proteins or lipids is due to a human lineage-specific genetic mutation in the enzyme cytidine monophosphate-N-acetylneuraminic acid hydroxylase (CMAH) ^22^. A direct consequence is that Neu5Gc epitopes are excluded from «self-tolerance» and elicit anti-Neu5Gc antibodies in humans ^17,23–26^.

Humans also lack the □1,3-galactosyl-transferase enzyme (GGTA1)^27^ and are not tolerant to □1,3 galactose (Gal) epitopes; they present various levels of natural anti-□1,3Gal antibodies and increase their level of anti-□1,3Gal antibodies after infusion of animal-derived products^28–30^.

Rabbit ATG given to diabetic patients resulted in highly significant increases in anti-Gal and - Neu5Gc IgG/IgM, still detectable one year post-infusion and responsible for the induction of SSD and immune complex disease in almost all patients^25^. In the context of concomitant administration of immunosuppressants and steroids, however, such as in organ transplantation, the administration of rabbit ALG/ATG rarely leads to SSD^11^. SSD, however, is a contributing factor to late graft loss following ALG/ATG induction^11^. High anti-Neu5Gc antibody level after kidney transplantation has also been associated with late graft loss^11^. This effect may be due to the well-known passive incorporation of Neu5Gc residues from food origin into endothelial and epithelial cells^31,32^, possibly resulting in a chronic inflammation called “xenosialitis” in the presence of anti-Neu5Gc antibodies, although the detrimental role of these “natural antibodies” in human still needs to be confirmed^24,33,34^. In parallel, anti-Gal antibody responses can be involved in anaphylaxis-type reactions targeting glycoconjugates present on therapeutic products of animal origin^23,25^.

To address these issues, LIS1, a “glyco-humanized” polyclonal serum, has been developed in pigs knocked out for the two genes CMAH and GGTA1 so that the two main xenogenic glycoantigens have been changed into human-type glycoantigens: the □1,3Gal epitopes are replaced by Gal β1,4-GlucNac, and Neu5Gc is replaced by Neu5Ac. They differ from rabbit immunoglobulins in humans since present the unique property to not interact with human FcγR, which modifies the mechanism of action by impairing ADCC^35^. Here, the mechanisms of action of LIS1 were investigated *in vitro* and *in vivo* in a nonhuman primates.

## 2. METHODS

### 2.1. LIS1 manufacturing and composition

Double KO defined high health status pigs were immunized with a human CD4^+^ CD8^+^ T-cell line. LIS1, the IgG fraction, was purified from serum in compliance with good manufacturing practice (GMP) and ICH guidelines. Details have been published elsewhere^35^..

### 2.2. Competition assay to determine the LIS1 target repertoire

Human peripheral blood mononuclear cells (PBMCs) were plated at 1.10^5^ cells per well in 96-well plates and then were labeled with LIS1 (250 μg/mL) and/or with the following monoclonal antibodies from BD Biosciences (Franklin Lakes, United States) at a 1/100 dilution: anti-human TCRa/b-FITC (Clone T10B9.1A.31); anti-human CD2-PE (Clone RPA-2.10); anti-human CD3-PE (Clone HIT3a); anti-human CD4-PE (Clone RPA-T4); anti-human CD8-PE (RPA-T8); and anti-human CD28-PE (CD28.2). A mean fluorescence intensity drop in flow cytometry indicated competition between LIS1 and the test antibody.

### 2.3. Determination of the LIS1 active fraction

The active fraction of LIS1 was determined by serial depletion against the cells used for immunization. Briefly, 30.10^6^ target cells fixed in 4% paraformaldehyde were resuspended in 1 mL of LIS1 (1.6 mg/ml) in phosphate buffered saline (PBS). After 20 min of incubation at room temperature, the cells were pelleted and the supernatant was transferred to a second tube containing a new fresh cell pellet to repeat the incubation. Five successive incubations were performed. The remaining specific antibodies were estimated/monitored after each incubation using a binding assay. The total IgG concentration was evaluated by spectrophotometry. The LIS1 active fraction was defined as the fraction of IgG cleared using this repeated depletion procedure.

### 2.4. Binding assay

PBMC, red blood cells and platelets from healthy donors were isolated in EDTA tubes. 10^5^ blood cells were incubated with increasing concentrations of pig or rabbit ALGs/ATGs for 30 min at 4 °C. After 3 washes, bound IgG was detected using FITC-conjugated anti-pig or anti-rabbit antibody (both from Bio-Rad) or AF-488 conjugated protein G (Thermo Fisher, Waltham, MA), and analyzed by flow cytometry.

### 2.5. Apoptosis assay

PBMCs were mixed with increasing doses of rabbit ATG or LIS1 in RPMI medium with 10% FCS. Nonimmune IgG was used as a negative control. After 3 h of culture (37 °C, 5% CO_2_), the cells were labeled with AF488-conjugated Annexin V and DAPI (Thermo Fisher) before analysis by flow cytometry. The percentages of cells in early apoptosis (Annexin V+/DAPI-cells) and late apoptosis (Annexin V+/DAPI+) were combined to determine the overall % of apoptosis.

### 2.6. Platelet aggregation assay

Human platelets were purified from citrated blood were mixed with increasing doses of LIS1 or rabbit ATG and aggregation was monitored for 60 minutes. To induce platelet aggregation, ristocetin (1.5 mg/ml), Thrombin Receptor Activator Peptide (TRAP,1.5 μM), arachidonic acid (0.3 mg/ml), ADP (adenosine diphosphate, 5 μM), epinephrine (5 μM) and collagen (1.25 μg/ml) were evaluated.

### 2.7. Opsonophagocytosis assay

Monocytes were purified from PBMCs by CD14 positive magnetic selection (Miltenyi Biotec, Bergisch Gladbach, Germany) and plated at 2.10^6^ cells/ml in RPMI 1640 medium containing 10% fetal calf serum (FCS). Monocyte differentiation into macrophages was performed for 5-7 days in the presence of human M-CSF (100 ng/ml) (R&D system). On the day of the assay, CFSE-labeled CD3 lymphocytes were incubated with ALG in RPMI medium with 10% heat-inactivated FCS for 30 min at 4 °C. Lymphocytes were washed twice with medium and cultured with human macrophages (ratio 1:1). After 3 hours of culture, the cells were washed twice, and macrophages were labeled with CD14-BV421 for 30 min at 4 °C, and analysis by flow cytometry. Phagocytosis was assessed as the percentage of double-positive (CFSE^+^/CD14^+^) cells over CD14^+^ cells.

### 2.8. Mixed Lymphocyte Reaction (MLR) assay

Proliferation was assessed using a CFSE dilution assay. Briefly, stimulator PBMCs were irradiated at 35 Gy and cultured (ratio 4:1) with CFSE-labeled responder PBMCs (Invitrogen, Waltham, Massachusetts, United States of America). The cells were cultured for 3 days in RPMI 10% FCS in the presence or absence of rabbit ATG or LIS1 and analysed by flow cytometry. Groups were compared using ANOVA and the Tukey□Kramer multiple comparison test. The results were expressed as %FITC low × number of lymphocytes to analyze cell proliferation of the residual cells.

### 2.9. *In vivo* evaluation

#### 2.9.1. Animals

Eighteen cynomolgus monkeys were used after authorization of the French Research Ministry (APAFIS 10717) / Citoxlab France Ethics Committee (CEC)/ Animal Welfare Body of Cynbiose/ Ethics Committee of VetAgro-Sup. The characteristics of these *in vivo* studies are reported in Supplementary Table 1. Cynomolgus monkeys received LIS1 by daily intravenous administration at doses ranging from 40 to 75 mg/kg/administration for 5 days. LIS1 was administered as a solution in the vehicle (sterile NaCl 0.9%) under a constant infusion rate.

**Table 1:** Antigenic targets. Human PBMCs were plated at 1.10^5^ cells per well. The cells were incubated with different labeled monoclonal antibodies (anti-human TCRα/β-FITC, anti-human CD2-PE, anti-human CD3-PE, anti-human CD4-PE, anti-human CD8-PE, anti-human CD28-PE) alone or in combination with LIS1 at 250 μg/mL. The residual labeling was analyzed by flow cytometry. The results are presented as the mean fluorescence intensity (MFI) and % signal inhibition by LIS1.

| Differentiation cluster (CD) | MFI |  | inhibition (%) |
| --- | --- | --- | --- |
|  | mAb | mAb+LIS1 |  |
| TCR$_{\alpha/\beta}$ | 1304 | 692 | 47 |
| CD2 | 4236 | 3262 | 23 |
| CD3 | 948 | 591 | 38 |
| CD4 | 2005 | 954 | 52 |
| CD8 | 1523 | 98 | 94 |
| CD28 | 535 | 400 | 25 |

#### 2.9.2. Lymphocyte depletion

CD3^+^ T lymphocyte numeration was performed using the BD Trucount system (BD Biosciences). Whole blood cells were labeled with the fluorochrome-labeled monoclonal antibody anti-CD3 (BD Biosciences). Other lymphocyte subpopulations were studied by flow cytometry. Briefly, fresh peripheral blood cells were stained with fluorochrome-labeled monoclonal antibodies against CD3 (SP34-2), CD4 (RPA-T4), CD8 (RPA-T8), CD20 (2H7), CD25 (M-A251), CD27 (T27.1), CD28 (CD28.2), CD31 (WM-59), CD45 (D058-1283), CD45-RA (5H9), CD95 (DX2), CD127 (hIL-7R-M21), GzmB, and FoxP3 (236A/E7) (Beckton Dickinson, San Jose, CA, USA). Polychromatic flow cytometric analyses were performed using a Canto II analyzer (BD Biosciences).

#### 2.9.3. Pharmacokinetic analysis

The LIS1 concentration in monkey serum was determined using pig IgG ELISA (Bethyl, Montgomery, Tx).

#### 2.9.4 Skin allograft model in cynomolgus monkeys

were used in the study. Skin grafts were performed on 5 anesthetized female cynomolgus monkeys (Supplemental Table 1). After back skin shearing and asepsis, the animals were grafted in pairs (back skin collection using a 20 mm diameter template, followed by grafting and skin graft suturing). In the back of each animal, three grafts were performed: one autograft and two pairwise allografts. In each group, one animal received skin transplants of his congener and vice versa. Three of them (Group 1) received LIS1 at 75 mg/kg once a day for 5 consecutive days. The injection was performed intravenously (concentration: 10 mg/mL; flow rate: 4 mL/kg/h; infusion duration: approximately 1 h50). The two remaining monkeys (Group 2) were used as controls and did not receive treatment.

### 2.10. Statistical analysis

All statistical analyses were performed using GraphPad Software (GraphPad Software, San Diego, CA). p values of <0.05 were considered statistically significant.

## 3. RESULTS

### 3.1. Specificity of target

#### 3.1.1. Binding of LIS1 to human blood cells

The binding of LIS1 to human PBMCs, platelets or red blood cells was investigated by flow cytometry (Figure 1A) and compared with that of rabbit ATG (Figure 1B). A dose-dependent and saturable signal was obtained. IgG from nonimmunized pigs used as negative control displayed negligible binding (Mean fluorescence intensity, MFI < 82). Binding to red blood cells was very low (MFI < 5000; Figure 1A) for LIS1 and rabbit ATG. Binding to platelets was barely detectable for both preparations. Since the comparison between LIS1 and rabbit ATG binding was questionable because of the two different detection systems (anti-pig and anti-rabbit antibodies), we used AF-488-conjugated protein G to similarly detect both pig- and rabbit-bound IgG. The ability of Protein-G to equivalently recognize pig and rabbit IgG was first confirmed by ELISA and surface plasmon resonance (data not shown). The binding of LIS1 to human platelets was significantly lower than that of rabbit ATG in the presence of Protein-G (Figure 1C). Next, we performed platelet assays to determine whether aggregation was altered or induced by LIS1 or rabbit ATG. No inhibition of induced platelet aggregation was observed with LIS1 (not shown). Both preparations induced platelet aggregation in this assay but more strongly and faster with rabbit ATG and at lower concentrations than with LIS1 (Figure 1D).

**Figure 1:**
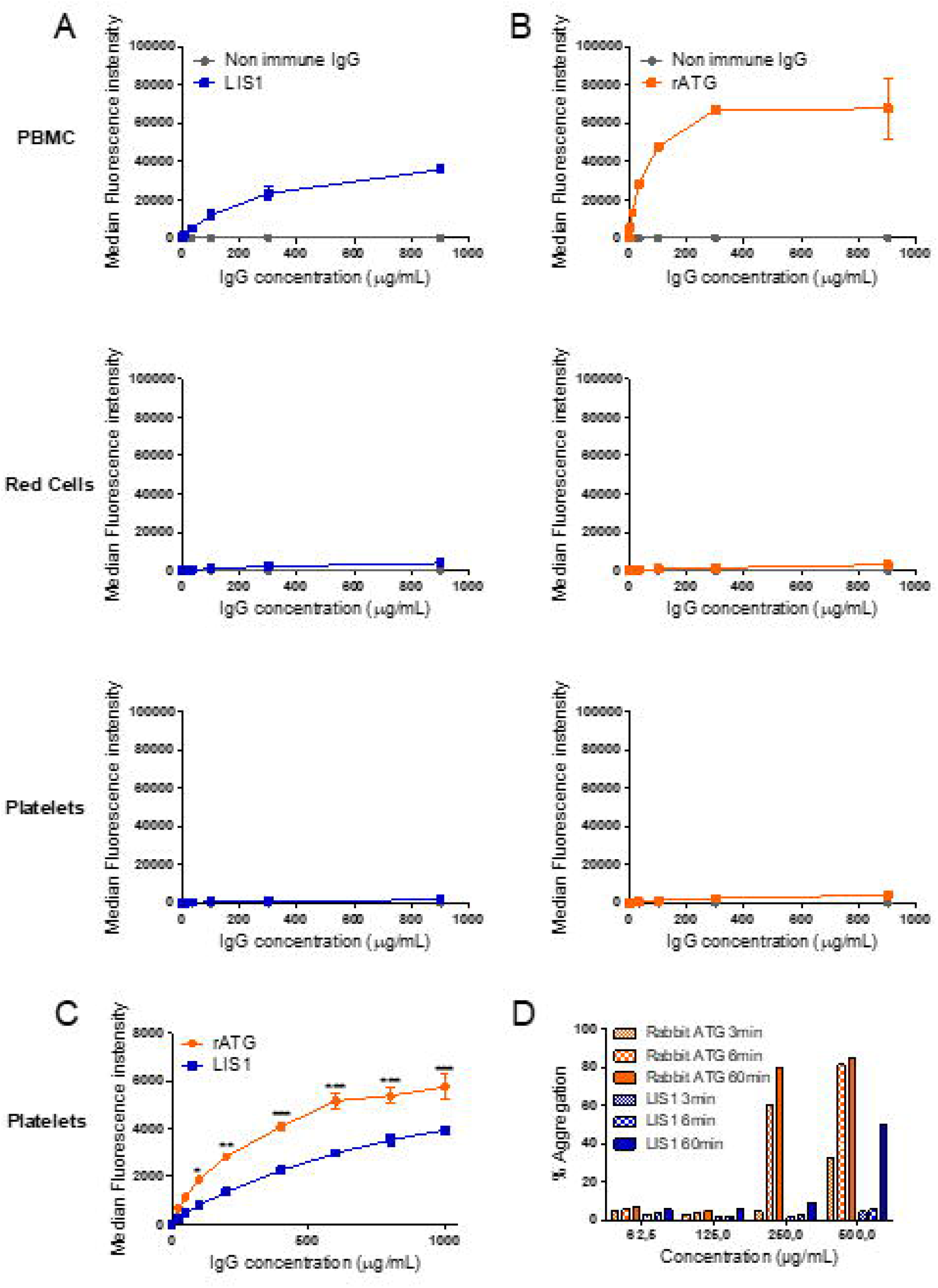
Interaction between LIS1 and human blood cells. Binding of LIS1 (**A)** or rabbit ATG (**B**) to human PBMCs, red cells and platelets was investigated by flow cytometry. Cells were incubated with increasing concentrations of LIS1 or rabbit ATG. Nonimmune IgG was used as a control. After washing, bound IgG was detected by flow cytometry using a FITC-conjugated anti-pig antibody or FITC-conjugated anti-rabbit antibody (N=3). **(C)** Direct comparison of LIS1 and rabbit ATG binding to human platelets. Human platelets were incubated with increasing doses of LIS1 or rabbit ATG. Attached antibodies were then detected using AF488-conjugated protein G and flow cytometry analysis. The data are expressed as means ± SEM (N=3). Two-way ANOVA (*, p<0.5; **, p<0.01; ***, p<0.005). **(D)** Platelet aggregation assays. Human platelets were purified from citrated blood, and aggregation was measured by turbidimetry using a TA-8 V platelet aggregometer (Stago). One representative experiment from two is shown.

#### 3.1.2 Antigenic targets and active fraction

Antigenic targets of LIS1 were determined using a competition assay against labeled-monoclonal antibodies targeting the classical T-cell markers—*i.e*., anti-TCR α/β, anti-CD2, anti-CD3, anti-CD4, anti-CD8 and anti-CD28 antibodies. In the presence of LIS1, we observed a strong decrease in the MFI for all antibodies evaluated (Table 1), indicating competition between mAbs and LIS1 on these targets.

We then determined the fraction of LIS1 actually binding target cells. Specific antibodies within LIS1 polyclonal IgG were removed by adsorption on human PBMCs (Supplementary Figure 1). The amount of nonspecific LIS1 antibodies remaining after 5 cycles of adsorption suggested that 40% of LIS1 IgG was specific to human PBMCs.

### 3.2. LIS1 mechanisms of action

ALG/ATG activity classically relies on lymphocyte depletion through complementary mechanisms such as CDC, ADCC, ADCP and apoptosis induction. We have previously shown that LIS1 is particularly efficient in recruiting human C1q and mediating CDC. By contrast, LIS1 could not interact with human Fcγ receptors and thus could not trigger ADCC in humans^35^. Here, we investigated other mechanisms of lymphocyte depletion using rabbit ATG as a reference to assess LIS1 activity.

#### 3.2.1. Apoptosis

Induction of apoptosis in human lymphocytes was investigated after exposure to increasing doses of LIS1. Nonimmune DKO pig IgG was used as a negative control. Induction of apoptosis, including early and late apoptosis, was monitored by flow cytometry using Annexin V and DAPI staining (Figure 2A). Significant apoptosis induction was detectable using 10 μg/ml of LIS1 (Figure 2B) and increased with LIS1 concentration, reaching a plateau at 300 μg/ml.

**Figure 2:**
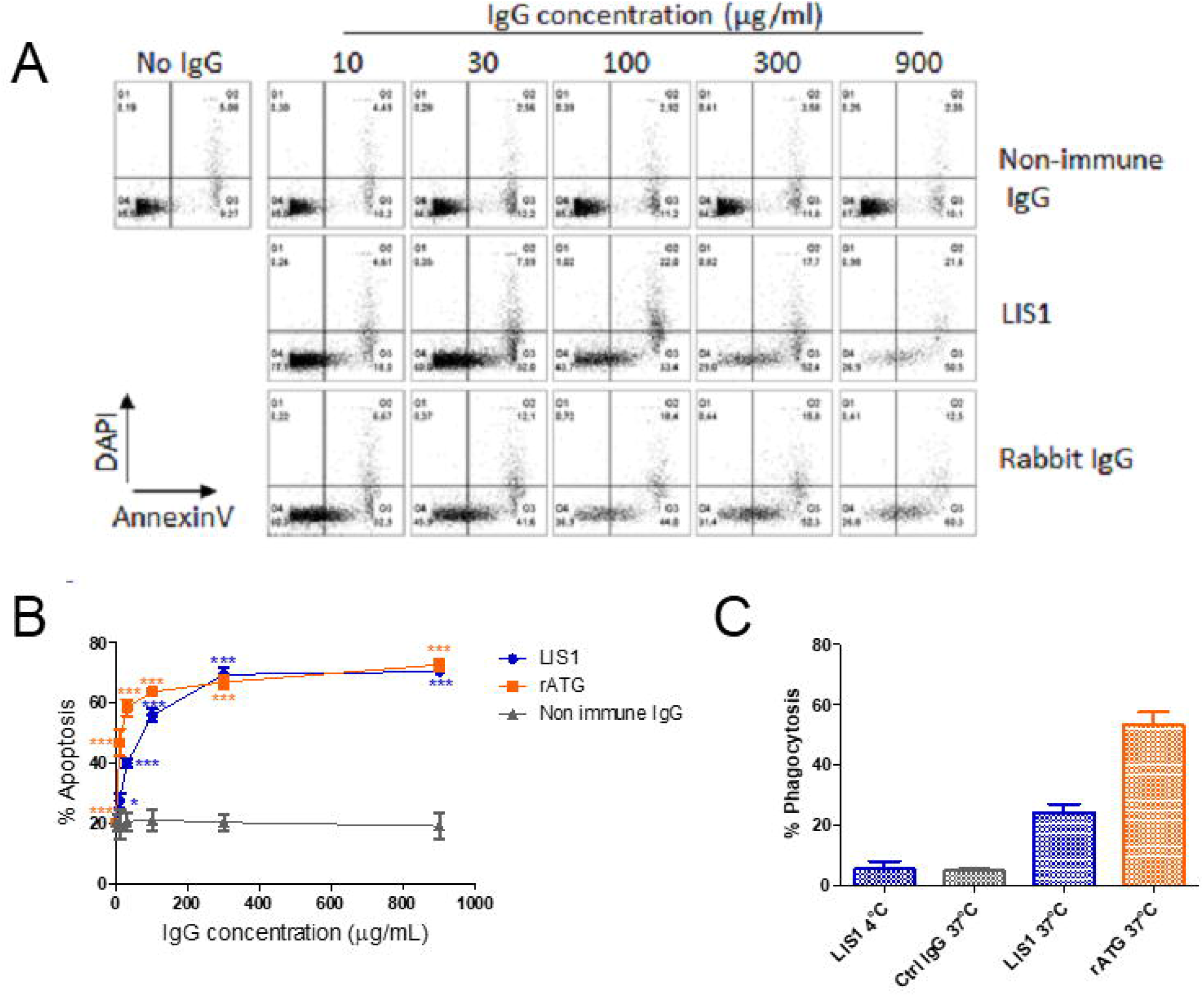
Mechanisms of action of LIS1. Induction of apoptosis by LIS1. **(A)(B)** Human PBMCs were treated with increasing doses (10, 30, 100, 300 and 900 μg/ml) of LIS1. Rabbit ATG and nonimmune DKO pig IgG were used as positive and negative controls, respectively. After 3 h of culture (37 °C, 5% CO_2_), apoptosis was monitored by flow cytometry by Annexin V and DAPI staining. (B) The mean percentage of apoptosis was then determined by adding the percentage of cells in early (Annexin V+/DAPI-) and late (Annexin V+/DAPI+) apoptosis. The data are expressed as means ± SEM (N=3). Two-way ANOVA (*, p<0.5; **, p<0.01; ***, p<0.005), comparison of LIS1 or rabbit ATG vs. nonimmune IgG. Phagocytosis assay. **(C)** Phagocytosis of opsonized PBMCs by human monocyte-derived macrophages was assessed by flow cytometry. CFSE-labeled cells were incubated with IgG in RPMI 10% FCS medium for 30 min at 4 °C. Lymphocytes were washed twice and cultured with human macrophages (ratio 1:1) in RPMI 10% FCS. After 3 hours of culture, the cells were washed twice, and macrophages were labeled with CD14-BV421 for 30 min at 4 °C. Cells were washed twice before flow cytometry analysis. Phagocytosis was assessed as the percentage of double[positive (CFSE+/CD14+) cells. Values were compared by ANOVA followed by Tukey’s post hoc test. ((*, p<0.5; **, p<0.01; ***, p<0.001).

#### 3.2.2. Phagocytosis of opsonized targets

Significant uptake of T lymphocytes by monocyte-derived macrophages was observed after opsonization with LIS1 or rabbit ATG (Figure 2C), with 23% or 53% of macrophages ingesting T cells, respectively. The absence of internalization when cultures were performed at 4 °C demonstrated the active process involved. The active LIS1 concentration was 10 μg/mL, a concentration with limited apoptosis. Therefore, the data presented reflected ADCP activity mediated by opsonization of targets and not phagocytosis of apoptotic cells.

#### 3.2.3 Inhibition of T-cell alloreactivity

We evaluated the ability of LIS1 to block T-cell alloreactivity. One-way MLRs were performed in the presence of LIS1 or rabbit ATG, and T-cell proliferation was monitored after 3 days. We observed a strong and dose-dependent inhibition of residual live T-cell proliferation when MLR was performed in the presence of LIS1 (Figure 3). The inhibition was 98% at a concentration of 500 μg/ml, contrasting with rabbit ATG which dose-dependently enhanced alloreactive T cells proliferation.

**Figure 3:**
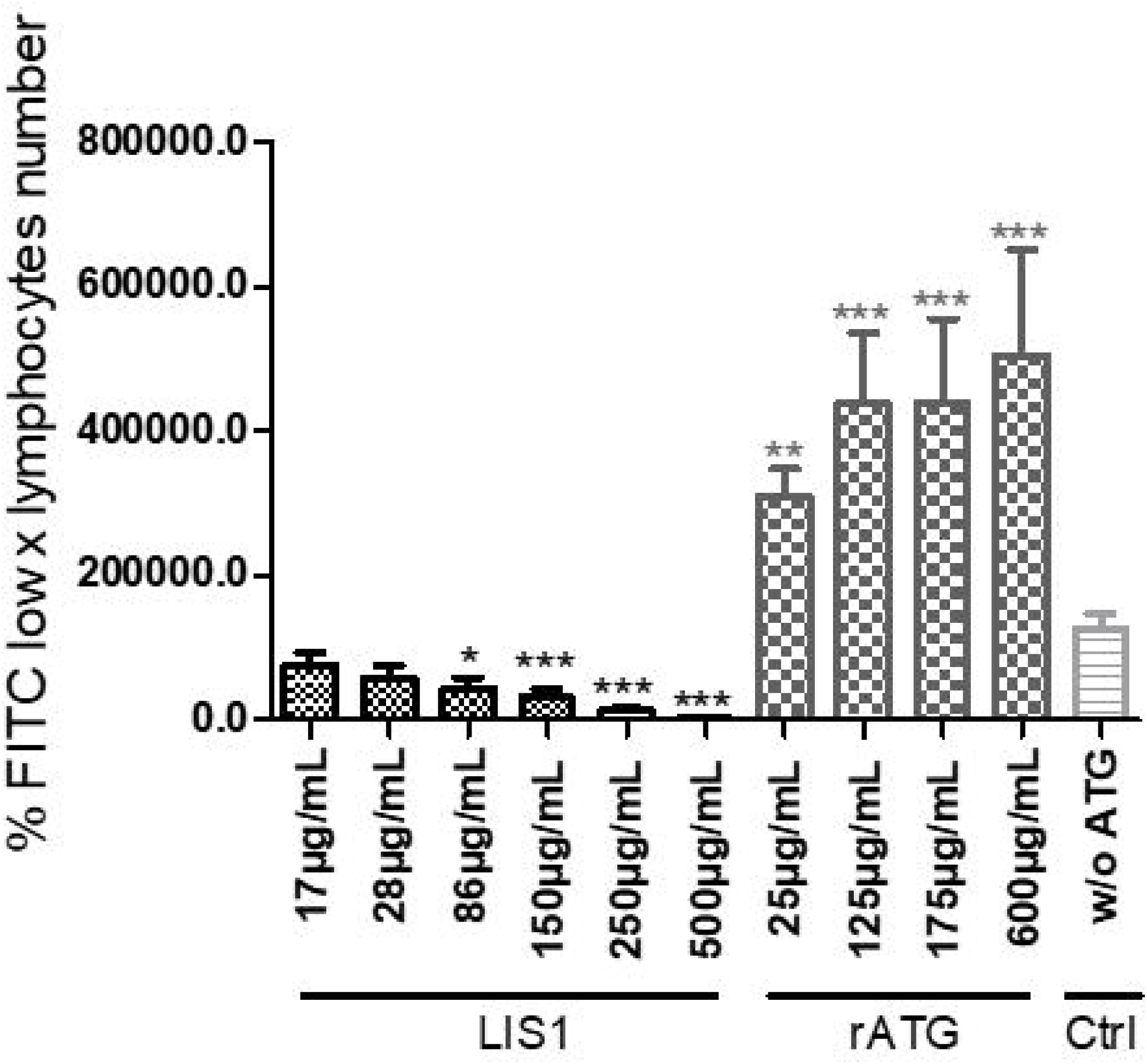
Mixed Lymphocyte Reaction assay. Stimulator human PBMCs were irradiated with 35 Gy; responder PBMCs were labeled with CFSE. Stimulator cells were adjusted to 8.10^6^ cells/mL, and responder cells were adjusted to 2 × 10^6^ cells/ml of RPMI medium. Next, the PBMCs were cocultured in a total volume of 200 μl of RPMI medium at 37 °C in a 5% CO_2_ incubator in the dark for 3 days with or without 250 μg/mL of rabbit ATG or LIS1. Proliferation was evaluated by measuring CFSE incorporation using flow cytometry. Groups were compared using ANOVA and the Tukey□Kramer multiple comparison test.

### 3.3. LIS1 in Nonhuman Primates

#### 3.3.1. Skin Graft rejection

To determine the ability of LIS1 to blunt alloreactivity *in vivo*, allogeneic skin grafts were performed in monkeys (Figure 4A). Graft rejection was defined by the presence of a scabby aspect, brown coloring with loss of flexibility, or complete necrosis of the epidermis. Rejection of allogeneic skin grafts occurred from Day 8 to 11 in the control group (untreated transplanted monkeys). Graft rejection was significantly delayed following induction treatment with LIS1 days 20-29 ^35^ (Figure 4B).

**Figure 4:**
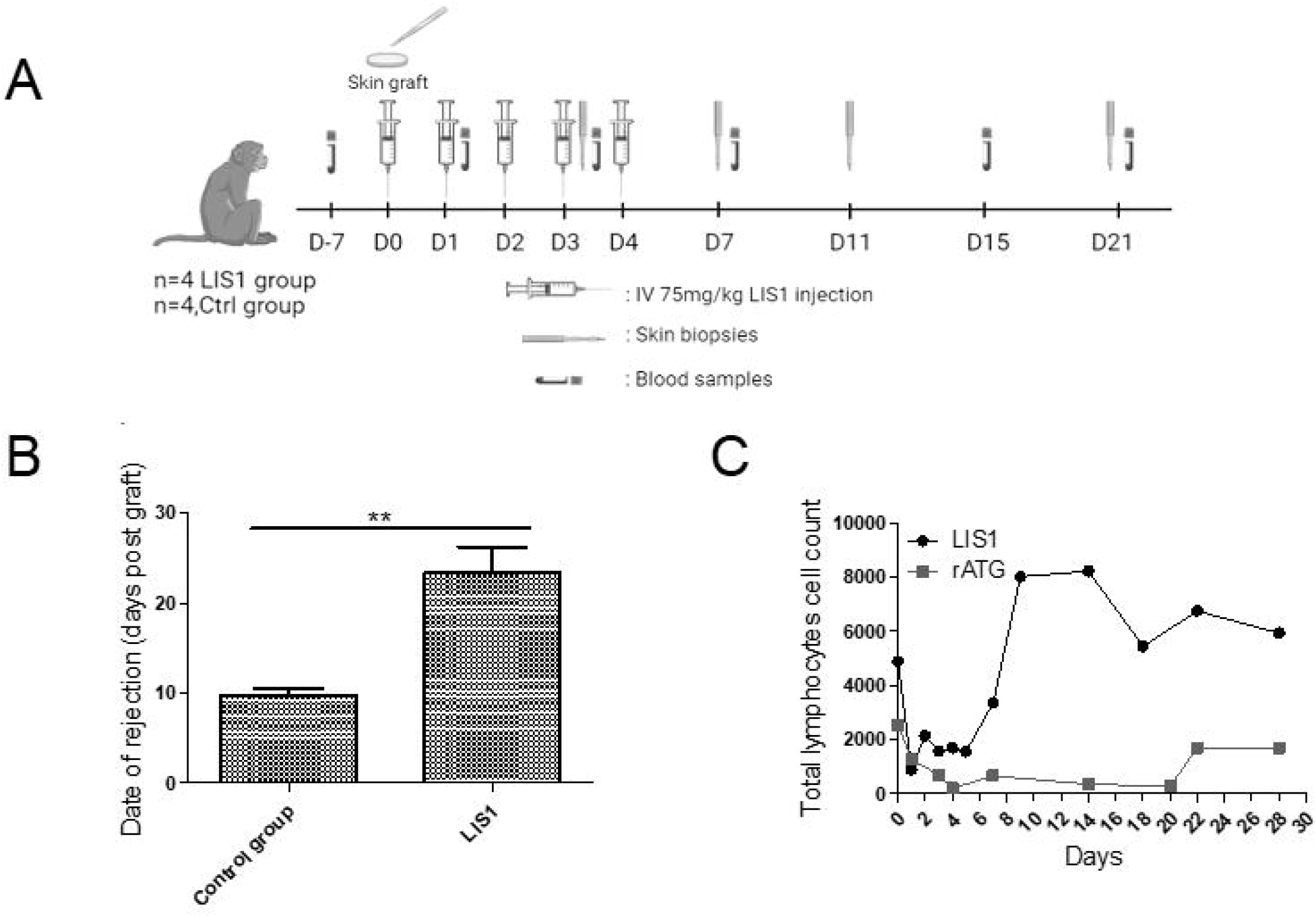
In vivo efficacy of anti-lymphocyte IgG in NHP. **(A)** Timeline of the NHP skin graft study. **(B)** Skin grafts were performed on anesthetized animals on Day 0. After back skin shearing and asepsis, the animals were grafted in pairs (back skin collection using a 20 mm diameter template, followed by grafting and skin graft suturing). In the back of each animal, three grafts were performed: one autograft (as a control) and two pairwise allografts (one animal received skin transplants of his congener). Four cynomolgus monkeys intravenously received LIS1 at 75 mg/kg once a day for 5 consecutive days. Four cynomolgus monkeys were used as controls and did not receive treatment. Skin graft rejection was considered when the area of the necrotized part covered the full graft, and the whole graft site hardened. **(C)** LIS1 and rabbit ATG-induced T-cell depletion in macaques. Cynomolgus monkeys (n=9) received I.V. daily doses of 50 mg/kg of LIS1 for 5 days. Cynomolgus monkeys (n=2) received I.V. daily doses of 5 mg/kg of rabbit ATG for 5 days.

#### 3.3.2. Total Lymphocyte depletion

Depletion experiments were also performed *in vivo* in cynomolgus monkeys. A dramatic drop in the CD3 lymphocyte count was observed after the first injection of LIS1 or rabbit ATG. (Figure 4C). However, lymphocyte depletion induced by LIS1 was limited to one week.

#### 3.3.3. Lymphocyte subpopulations

We investigated the depletion of lymphocyte subpopulations after five I.V. infusions of LIS1 (50 mg/kg/adm) in cynomolgus monkeys. LIS1 induced selective depletion of peripheral effector naive T cells (CD4^+^ TN p=0.005; CD8^+^ TN, p=0.0002), memory effector T cells (CD4^+^ EM p=0.0134; CD8^+^ EM p=0.002) and recent thymic embryonic cells (CD4^+^ RTE, p=0.007; CD8^+^ RTE, p<0.007) between Day 0 and Day 3/4 (Figure 5). Depletion was stronger for CD8 cells (86% drop for naive CD8 cells and 95% for memory effectors) than CD4 cells (71% decrease for naive CD4 cells and 43% for memory effectors). By contrast, B cells and Tregs were unaffected by LIS1 (Figure 5).

**Figure 5:**
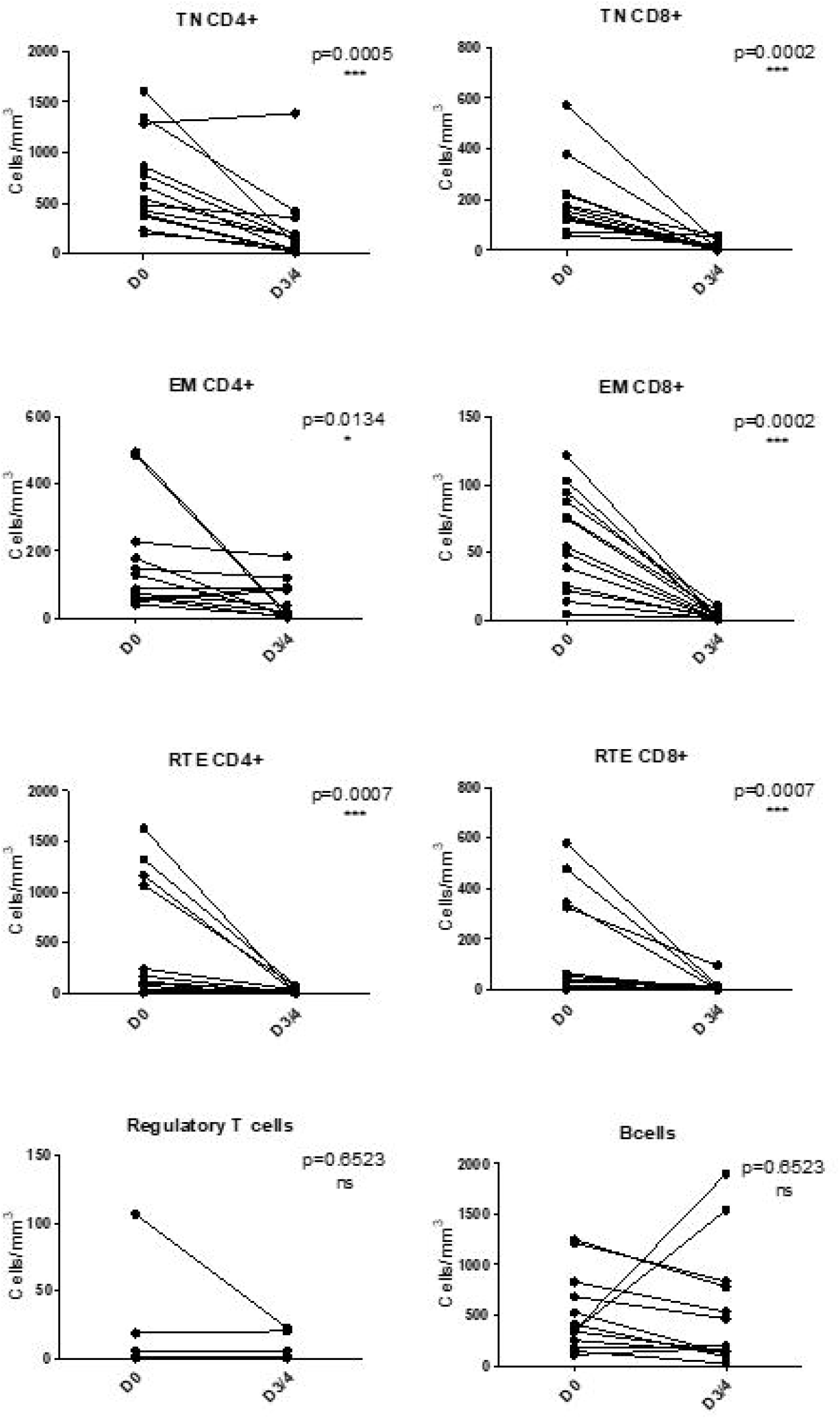
LIS1 shows selective immunoreactivity to peripheral T cells. Cynomolgus monkeys (n=11) received I.V. daily doses of 50 mg/kg LIS1 for 5 days. On Days 3-4, blood was sampled, and lymphocyte subpopulations were analyzed by flow cytometry. Whereas CD4+ and CD8+ T-cell subpopulations were reduced, Treg and B cells were unaffected. Monocytes were also unaffected (not shown).

#### 3.3.4. Pharmacokinetic study

Serum samples were drawn and analyzed by ELISA. LIS1 presented a T_1/2_ half-life of approx. 40 h, and a distribution profile compatible with the plasmatic compartment in primates (Figure 6). The mean systemic clearance was 0.044 and 0.048 mL/h/kg in females and males, respectively. Quantifiable serum concentrations were observed until the last collected time point (29 days after the last infusion on Day 5), reflecting a slow elimination rate of LIS1. No marked difference was observed between males and females, as the ratio of AUC_0-t_ values was close to 1.

**Figure 6:**
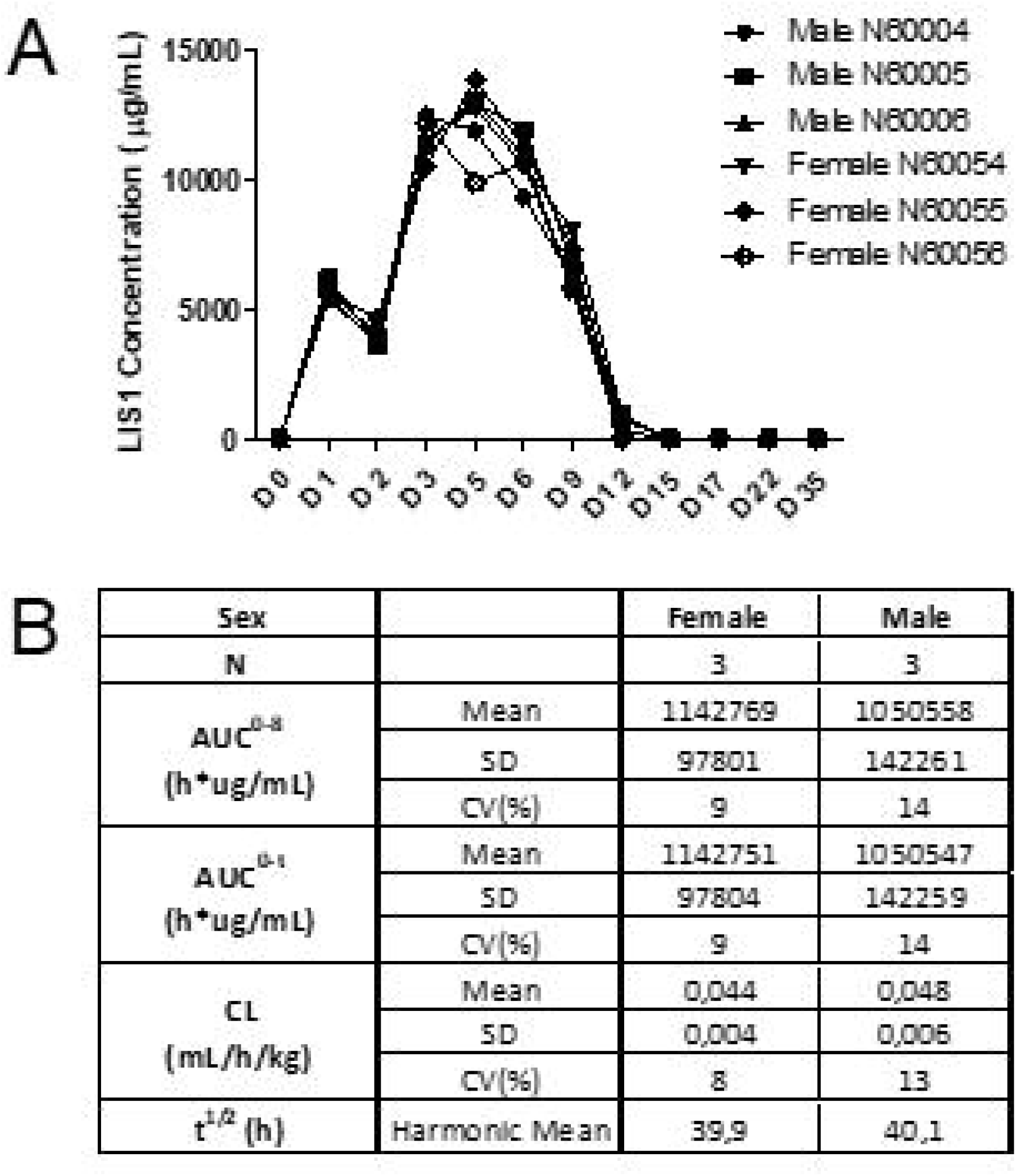
Pharmacokinetic evaluation. For the LIS1 GLP toxicology study, one group of three male and three female cynomolgus monkeys received LIS1 by daily intravenous administration (4-hour infusion) at 50 mg/kg/adm for 5 days. Serum samples were drawn on Days 1 (before infusion, 5 minutes, 24 and 96 hours after infusion) and 5 (before infusion and 5 minutes after infusion) and Days 6, 9, 12, 15, 17, and 22 and week 5. Pharmacokinetic parameters were analyzed by ELISA. (**A)** Pharmacokinetic profiles of LIS1-treated macaques. **(B)** Pharmacokinetic characteristics.

## 4. DISCUSSION

LIS1 is a novel ALS obtained from GGTA1 and CMAH double KO pigs hyperimmunized with a double-positive human CD4^+^/CD8^+^ T-cell line. LIS1 differs from other ALSs by its mechanism of action, which relies on apoptosis, CDC and inhibition of alloreactivity without ADCC.

The goal of induction therapy is to control early acute rejection following organ transplantation^36,37^. After infusion, rabbit ATG induces immediate immune cell depletion, particularly T lymphocyte depletion, through ADCC, CDC, opsonization resulting in phagocytosis of IgG-coated T cells apoptosis by activation of the Fas-Fas ligand pathway ^38–40^. Although these mechanisms allow marked depletion of target cells, they come at the cost of leukopenia and thrombocytopenia. Moreover, in some patients, CD4^+^ lymphopenia lasts for several months, increasing the risk of infection and other complications^41,42^.

One main challenge in organ transplantation is to define the best immunosuppressive strategy to ensure long-term graft and patient survival by preventing graft rejection and protecting against treatment side effects^2,41,42^. Thrombocytopenia is a frequent complication, partly induced by immunosuppressive therapies. Thrombocytopenia is more common in patients receiving ATGs than in those receiving basiliximab in the first few days after transplantation^2,42^. Although thrombocytopenia is often mild and transient, for kidney transplantation, it may exacerbate the perioperative coagulation imbalance in patients with already compromised bleeding control. Because of differences in the source of cells used for immunization, LIS1 and other ATGs have different antigenic profiles, and these differences may lead to clinical consequences^43^, particularly for thrombocytopenia. We illustrate here that the binding and platelet aggregation of LIS1 is very low, which might result in lower thrombocytopenia.

The mode of action of LIS1 includes transient immunosuppression by inducing CDC, ADCP and apoptosis-mediated selective T-cell depletion. Using *in vitro* assays and *in vivo* pharmacodynamic and transplantation models in nonhuman primates (NHPs) or humanized RRGS rats^44^, LIS1 confirmed this mode of action and nevertheless demonstrated a strong impact on T-cell responses in a stringent skin graft rejection model.

Conventionally, ADCP is described as a mechanism by which antibody-opsonized target cells activate FcγRs on the surface of macrophages to induce phagocytosis^45^. Thus, the mechanism underlying LIS1-generated ADCP is currently unknown because no interaction exists between porcine IgG and human FcγRs, except for FcRn^35^. Studies have shown that FcRn expression on phagocytic cells^46,47^ increases the phagocytosis capacity of opsonized IgG particles^48,49^. This mechanism of phagocytosis^49,50^ would likely require external sensing and cellular activation via FcγR or pattern recognition receptors (C-type lectin receptors)^51^. Phagocytosis by neutrophils and monocytes mediated by the interaction between C3b fragment from the cleavage of complement protein C3 and the CR1 receptor (CD35) is also a possible mechanism, although not explored here.

In a humanized mouse model ^52^, rabbit ATG has been shown to bind almost all human hematopoietic cell lineages, including hematopoietic stem cells, and thus likely depletes all subpopulations. The prolonged lymphodepletion observed in macaques treated with rabbit ATG, where dose-dependent depletion of CD3^+^ T cells is maintained and T-cell count recovery is delayed to several weeks^53^, might originate from this broad impact of ATG on T cells and stem cells, also impacting T-cell reconstitution. Because LIS1 is obtained by immunization using a selected effector-type human T-lymphocyte cell line, it is likely to show narrower immunoreactivity toward T lymphocytes than rabbit ATG obtained from rabbits immunized with a preparation of human thymocytes containing mature and immature T cells, Treg cells, progenitor cells, dendritic cells, epithelial cells and endothelial cells. Consequently, LIS1 shows selective immunoreactivity to peripheral effector T cells but not to B cells Treg cells, recent thymic emigrant cells and myeloid-derived suppressor cells described in experimental models and humans to contribute to immunosuppression maintenance post kidney transplantation^54,55^. *In vivo* evaluation in nonhuman primates showed that LIS1 robustly depleted target T cells from the peripheral blood. Interestingly, depletion was intense but short-term because T-cell counts recovered within two weeks. In a previous study in RRGS (humanized) rats reconstituted with human PBMCs^44^, LIS1 blocked human T-cell reconstitution by more than 98% and prevented graft versus host disease (GVHD).

Additionally, *in vitro*, LIS1 demonstrated potent immunosuppressive activity in MLR experiments. As shown in Table 1 LIS1 recognized and presumably blocked T-cell receptors crucial for alloreactivity, without intrinsic agonist activity. This feature might be due to the incompatibility between LIS1 and human Fc receptors, reducing FcγR-mediated cross-linking and agonist events. This mechanism is similar to that of another class of induction therapy, anti-CD25 monoclonal antibodies (e.g., basiliximab), which completely and consistently block activated T cells and their proliferation by blocking the alpha chain of the interleukin-2 receptor.

Although potent T-cell depletion has been historically considered necessary to prevent acute rejection during the early posttransplant phase for patients with high immunological risk, the prolonged T-cell depletion lasting several weeks observed with current ATG/ALG therapies^38^ impaired CD4^+^ T-cell reconstitution ^56^ are associated with an increased risk of infection, including bacterial infections (urinary tract and wound infections) and less common viral infections, such as cytomegalovirus (CMV), Epstein□Barr virus (EBV) and BK polyomavirus (BKV) infections^57^. Studies have also reported that CD4^+^ T-cell lymphopenia is associated with infectious complications^58^, cardiovascular complications ^59^ and mortality. An increased risk of infections is associated with the increased risk of posttransplant morbidity, rehospitalization and overall mortality^60,61^ and with decreased graft survival due to the infection itself and to the reduction in immunosuppressive therapy needed to manage infections^62^. Our preclinical data show that after 5 consecutive days of LIS1 infusion, T-cell depletion occurs acutely over 5 days and returns to baseline levels at the latest within 3 weeks (see Figure 4). If confirmed in human, these data suggest that LIS1 may reduce the risk of infection associated with prolonged depletion observed with other ALS therapies. This regimen would for example benefit patients with a higher risk of infection, among which elderly kidney graft recipients^62^. Avoiding prolonged T-cell depletion may also reduce the risk of neoplasm in patients, a possible additional benefit from induction with LIS1^63^.

LIS1 presented shorter and milder depleting activity than rabbit ATG and exhibited direct T-cell blockade activity in a mixed lymphocyte reaction. Whether the net effect of these differences allows similar protection against allograft rejection still remains to be proven in future clinical studies. *In vivo* efficacy data in a stringent skin graft model in cynomolgus monkeys showed an equivalent capacity of LIS1 to prevent allograft rejection compared to published Thymoglobulin® data^53^, 9 to 20-25 days with Thymoglobulin® and 9 to 20-29 days with LIS1.

Breg cells prevent graft rejection and belong to the signature of tolerance ^64–66^. Our study shows that LIS1 spares B cells, thus avoiding depletion of B-regulators, a major difference from rabbit ATG, which induces apoptosis of all B-cell subsets^67^. Similarly, regulatory T cells (Tregs) play a crucial role in limiting renal transplant rejection and, potentially, in promoting transplant tolerance^68^. Rabbit ATG induces and expands CD4^+^CD25^+^Foxp3+ cells with regulatory function ex vivo^69^. We observed that Tregs were not affected by LIS1 in nonhuman primates, at least in the short term. Whether these differences in the modulation of regulatory B and T cells may impact the outcomes of induction treatment is unknown.

LIS1 is currently undergoing clinical evaluation in humans in de novo kidney transplant recipients (ClinicalTrials.gov Identifier: NCT04431219). This trial will investigate whether the short-term depletion observed in primates replicates in humans and allows safe induction treatment. The hypothesis of an improved posttransplant outcome due to faster T-cell reconstitution and avoidance of infections, while similarly controlling acute rejection needs to be assessed in larger cohorts.

Our paper suffers from limitations related to the complexity of the mode of action of polyclonal products and of the induction procedure in solid organ transplantation. Ideal duration of lymphodepletion post-induction is unclear, and it is unknown whether the short-term T cell depletion reported with LIS1 plus the LIS1-induced blockade of alloreactivity will be sufficient to blunt acute rejection to the same extent as long-term depleting ATG. Although absence of B cell depletion by LIS1 might preserve Breg cells important in tolerance induction, it is unknown whether this presents advantage or drawback, owing to the possible lower control of anti-donor antibodies. Finally, the potentially different active titers of specific antibodies make elusive quantitative comparison of LIS1 and rabbit ATG. However, our data suggest that LIS1 may gather a specific pattern of properties that could increase the clinical advantage/disadvantage ratio of polyclonal anti human T cell preparations and widen their clinical usage.

In summary, LIS1 is a novel induction agent with potential interest for transplanted patients, overcoming problems related to xeno-antigenicity and prolonged target T-cell depletion. Its mechanism of action, combining mild T-cell depletion with inhibition of alloreactivity, warrants its clinical evaluation in kidney graft recipients.

## Abbreviations

ALG: anti-lymphocyte serum
aGal: alpha galactose
ADCC: antibody-dependent cell-mediated cytotoxicity
AUCinf: area under the curve ad infinitum
PK: pharmacokinetic
Neu5Gc.: 

## Authorship

Conceived the study: JMB, SB, OD, JPS, BV

Designed and supervised some experiments: OD, JPS, BV

Performed the experiments: CC, GE, JR, PJR, AS

## Acknowledgments

The authors thank Marc Fouassier (CHU Nantes) for his help in the platelet aggregation assay.

## Data Availability Statement

The data that support the findings of this study are available from the corresponding author upon reasonable request.

## FIGURE LEGENDS

***<u>Supplemental Table 1:</u> Overview of the in vivo analysis of cynomolgus monkeys***. *Cynomolgus monkeys (n=16) received I.V. daily doses ranging from 40 to 75 mg/kg LIS1 for 5 days. The characteristics of these in vivo studies are described here*.

***<u>Supplemental Figure 1</u>***: ***Evaluation of the LIS1 active fraction***. *One milliliter of LIS1 diluted in PBS to a concentration of 1.6 mg/mL was incubated with 30.10*^*6*^ *target lymphocyte cells. After 20 min of incubation at room temperature, the supernatant (SN1) was transferred to a second tube containing 30.10*^*6*^ *target T lymphocytes. After 20 min of incubation at room temperature, the supernatant (SN2) was collected and stored until SN5. The supernatants obtained were serially diluted and then incubated (30 min, 4 °C) with fresh XT1501 cells. After washing, a secondary anti-pig antibody was deposited, revealing the remaining XT1501-specific antibody fraction. LIS1 was used as a positive specificity control. Nonimmune IgGs were used as a negative control*.

