## Supplementary figures and images for "LIS1, a glyco-humanized swine polyclonal anti-lymphocyte globulin, as a novel induction treatment in solid organ transplantation"

### Supplemental Figure 1

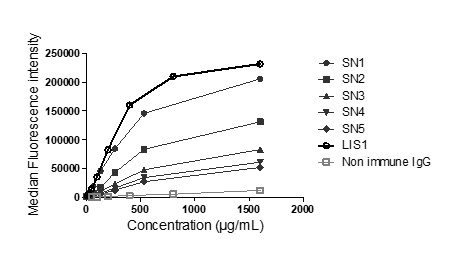

### Supplemental Table 1

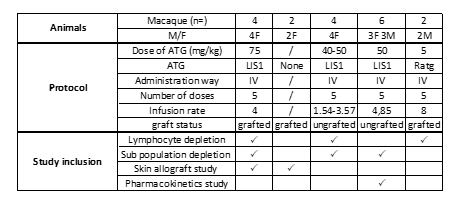
